# *Listeria monocytogenes* exploits the MICOS complex subunit Mic10 to promote mitochondrial fragmentation and cellular infection

**DOI:** 10.1101/712067

**Authors:** Filipe Carvalho, Anna Spier, Thibault Chaze, Mariette Matondo, Pascale Cossart, Fabrizia Stavru

## Abstract

Mitochondrial function adapts to cellular demands and is affected by the ability of the organelle to undergo fusion and fission in response to physiological and non-physiological cues. We previously showed that infection with the human bacterial pathogen *Listeria monocytogenes* elicits transient mitochondrial fission and a drop in mitochondrial-dependent energy production through a mechanism requiring the bacterial pore-forming toxin listeriolysin O (LLO). Here, we performed quantitative mitochondrial proteomics to search for host factors involved in *L. monocytogenes*-induced mitochondrial fission. We found that Mic10, a critical component of the mitochondrial contact site and cristae organizing system (MICOS) complex, is significantly enriched in mitochondria isolated from cells infected with wild-type but not with LLO-deficient *L. monocytogenes*. Increased mitochondrial Mic10 levels did not correlate with upregulated transcription, suggesting a post-transcriptional regulation. We showed that Mic10 is necessary for *L. monocytogenes*-induced mitochondrial network fragmentation, and that it contributes to *L. monocytogenes* cellular infection independently of MICOS proteins Mic13, Mic26 and Mic27. Together, *L. monocytogenes* infection allowed us to uncover a role for Mic10 in mitochondrial fission.

**Importance:** Pathogenic bacteria can target host cell organelles to take control of key cellular processes and promote their intracellular survival, growth, and persistence. Mitochondria are essential, highly dynamic organelles with pivotal roles in a wide variety of cell functions. Mitochondrial dynamics and function are intimately linked. Our previous research showed that *Listeria monocytogenes* infection impairs mitochondrial function and triggers fission of the mitochondrial network at an early infection stage, in a process that is independent of the main mitochondrial fission protein Drp1. Here, we analyzed how mitochondrial proteins change in response to *L. monocytogenes* infection and found that infection raises the levels of Mic10, a mitochondrial inner membrane protein involved in formation of cristae. We show that Mic10 is important for *L. monocytogenes*-dependent mitochondrial fission and infection of host cells. Our findings thus offer new insight into the mechanisms used by *L. monocytogenes* to hijack mitochondria to optimize host infection.

## Introduction

Mitochondria constitute one of the most important eukaryotic organelles due to their role in several essential cellular processes, such as energy production, biosynthesis of metabolic intermediates, calcium storage and signaling, autophagy, apoptosis, as well as redox and innate immune signaling (1–3). The overall morphology and cellular distribution of the mitochondrial network are controlled by a succession of fusion and fission events referred to as “mitochondrial dynamics”. This dynamic equilibrium is fundamental to meet cellular energetic and metabolic demands and respond to stress-inducing conditions (4).

Mitochondrial dynamics are governed by a family of large GTPases with membrane-shaping properties necessary to drive fusion or fission of mitochondria (5, 6). Mitochondrial fusion requires the sequential merging of the outer (OMM) and inner mitochondrial membranes (IMM) by the action of the OMM-anchored mitofusin 1 (Mfn1) and 2 (Mfn2) proteins, and the IMM-bound optic atrophy 1 (Opa1) protein. Mitochondrial fission is mainly mediated by the cytosolic dynamin-related protein 1 (Drp1), although the endoplasmic reticulum, actin and septins also play significant roles (6, 7). The importance of mitochondrial dynamics in cell physiology is attested by numerous neuro-muscular pathologies associated with genetic defects affecting the expression or activity of proteins involved in mitochondrial fusion and fission (8).

Due to their involvement in essential cellular processes, mitochondria are an attractive target for viral and bacterial pathogens (9–12). Indeed, many pathogenic bacteria were shown to modulate mitochondrial dynamics to create the ideal conditions for intracellular replication, immune evasion and persistence (11, 13–17). We previously explored how mitochondrial function and dynamics are impacted by infection with *Listeria monocytogenes* (16, 18), a facultative intracellular bacterial pathogen responsible for listeriosis, a life-threatening disease in immunocompromised individuals (19). We showed that *L. monocytogenes* causes fragmentation of the host mitochondrial network early in infection. This event requires the bacterial pore-forming toxin listeriolysin O (LLO), which promotes calcium influx into the host cell (16), causing a drop of the mitochondrial membrane potential and Drp1-independent mitochondrial fission (18). *L. monocytogenes* infection has thus revealed an unconventional mechanism of mitochondrial fission, but the mechanistic details and molecular players involved in modulation of mitochondrial dynamics and function upon *L. monocytogenes* infection still remain unclear.

Here, we set out to increase our understanding of the impact of *L. monocytogenes* infection on host cell mitochondria and identify novel factors involved in *L. monocytogenes*-induced mitochondrial fission by performing a quantitative characterization of the mitochondrial proteome upon infection. We report that *L. monocytogenes* infection significantly upregulates the mitochondrial levels of Mic10, a core subunit of the mitochondrial contact site and cristae organizing system (MICOS) complex (20). We show that this increase in Mic10 abundance requires LLO but is not correlated with increased transcription. Finally, we demonstrate that Mic10 is necessary for *L. monocytogenes*-induced mitochondrial fragmentation, and contributes to bacterial infection.

## Results

### Quantitative proteomic analysis of the human mitochondrial response to *L. monocytogenes* infection

To understand how the human mitochondrial proteome is affected by *L. monocytogenes* infection, we performed quantitative, label-free proteomic analysis of mitochondria isolated from human cells infected with *L. monocytogenes* (Figure 1A). We used HCT116 cells, a human intestinal epithelial cell line rich in mitochondria and efficiently infected by *L. monocytogenes*. Since *L. monocytogenes*-induced mitochondrial fission occurs early in infection and requires LLO (16), we performed short infections (2 h) with wild-type *L. monocytogenes* or with an LLO-deficient strain to focus on LLO-dependent processes. Mitochondria were isolated from infected and uninfected cell lysates by magnetic immunoaffinity purification and mitochondria-associated proteins were processed for LC-MS/MS analysis (Figure 1A). A total of 2,370 unique proteins were identified, with 2,039 (86%) proteins detected in every condition (Figure 1B). Among all identified proteins, 862 (36.4%) were annotated as mitochondrial (Figure 1B), which represents a good mitochondrial enrichment degree in our samples (compared to 7–8% of mitochondrial proteins in the human proteome) and a high coverage of the annotated mitochondrial proteome (53% of 1626 proteins; IMPI version Q2, June 2018). This overrepresentation of mitochondrial proteins is reflected in the results of a Gene Ontology (GO) term enrichment analysis, showing eight mitochondrial terms among the ten most enriched GO biological processes (Figure 1C).

**Figure 1 –.**
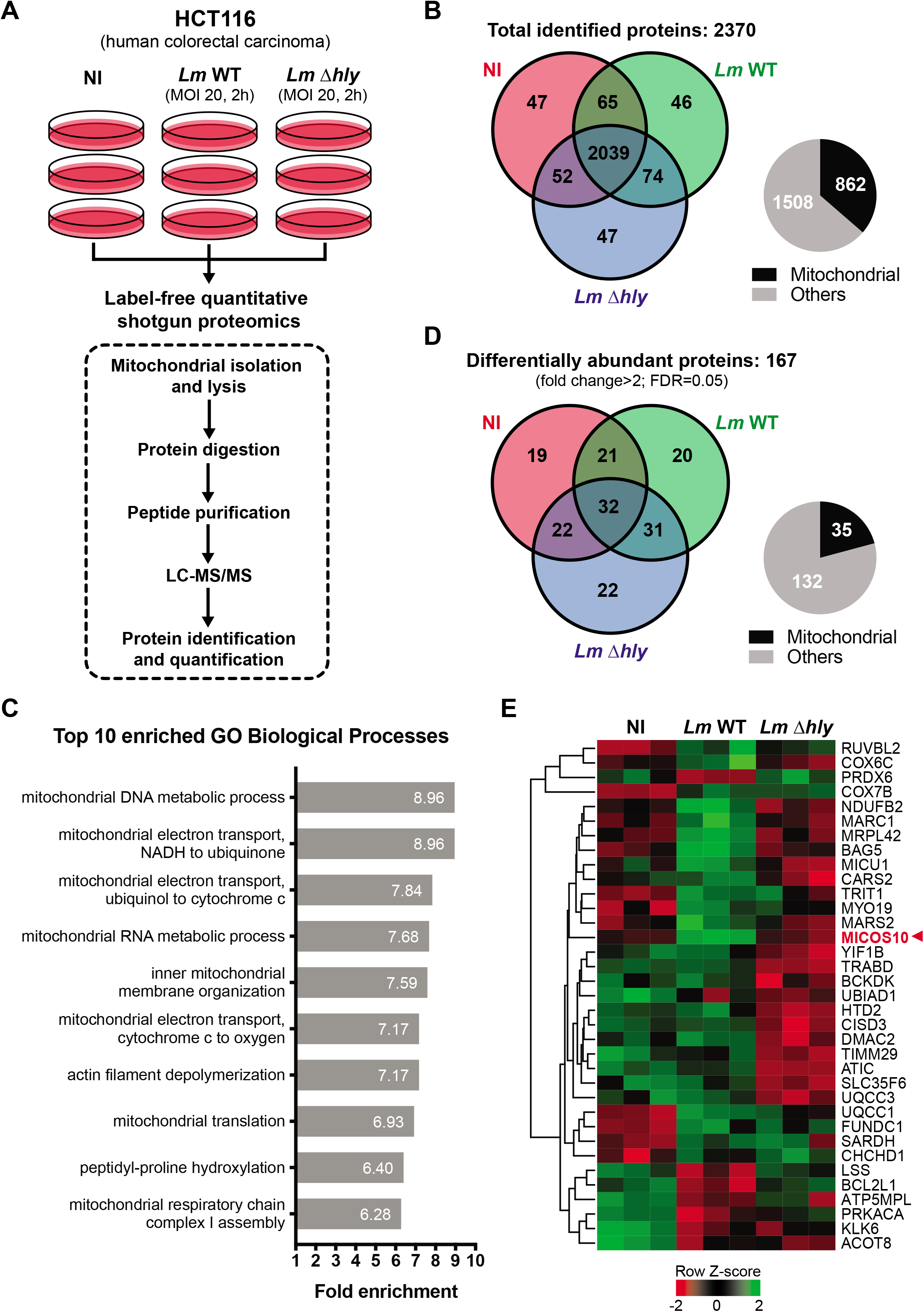
Analysis of changes in the human mitochondrial proteome elicited by *L. monocytogenes* infection. A) Schematic diagram of the experimental procedure used for proteomic analysis of human mitochondria isolated from cells infected or not with *L. monocytogenes*. B) Venn diagram of unique and common proteins identified in mitochondria isolated from uninfected and *L. monocytogenes-infected* cells. Pie chart shows distribution of proteins classified as mitochondrial and non-mitochondrial according to MitoMiner’s IMPI database (Q2, 2018). C) Top ten most prevalent Gene Ontology (GO) biological processes obtained from functional enrichment analysis (PANTHER gene analysis tool) of all proteins identified in mitochondria isolated from uninfected and *L. monocytogenes-infected* cells. D) Venn diagram of unique and common proteins identified in mitochondria isolated from uninfected and *L. monocytogenes*-infected cells with statistically significant changes in their abundance (fold change>2, FDR=0.05). Pie chart shows distribution of differentially abundant proteins classified as mitochondrial and non-mitochondrial according to MitoMiner’s IMPI database (Q2, 2018). E) Heat map showing relative changes in abundance of the 35 proteins annotated as mitochondrial in D. For each protein, the intensity levels in every condition (each box represents a triplicate) were normalized to the mean value using Z-score. Intensity levels higher and lower than the mean are indicated in green and red shades, respectively. Proteins are indicated by their gene symbol.

To determine proteins displaying significantly altered levels with infection, we performed a statistical test on all identified proteins to select those whose abundance changed at least two-fold. We obtained a list of 167 proteins, of which 32 (19.1%) were detected in every condition (Figure 1D). To find proteins that could play a role in mitochondrial fission, we focused on *bona fide* mitochondrial proteins or predicted to be functionally associated and/or interact with mitochondria. According to the IMPI, 35 of the 167 differentially abundant proteins (21%) were annotated as mitochondrial (Figure 1D, Table S1). Among these are proteins involved in the mitochondrial electron transport chain, such as the NADH:ubiquinone oxidoreductase subunit B2 (NDUFB2) and assembly factor DMAC2, the ubiquinol-cytochrome c reductase assembly factors 1 and 3 (UQCC1 and UQCC3), the cytochrome c oxidase subunits 6C and 7B (COX6C and COX7B), and the F_1_F_O_ ATP synthase subunit 6.8PL (ATP5MPL). Four of these proteins become significantly more abundant in response to infection, suggesting an increased activity of the respiratory chain. Other differentially abundant proteins identified in our analysis are associated with mitochondrial translation (cysteinyl- and methionyl-tRNA synthetases 2, CARS2 and MARS2; tRNA isopentenyltransferase 1, TRIT1; and mitochondrial ribosomal proteins CHCHD1 and MRPL42), metabolism of sterols (lanosterol synthase, LSS), fatty acids (hydroxyacyl-thioester dehydratase type 2, HTD2) and branched-chain amino acids (branched-chain ketoacid dehydrogenase kinase, BCKDK), regulation of mitophagy (Bcl2-associated athanogene 5, BAG5 (21, 22); FUN14 domain-containing 1, FUNDC1 (23); peroxiredoxin 6, PRDX6 (24)) and apoptosis (Bcl2-like protein 1, BCL2L1), cristae formation (MICOS complex subunit Mic10, MICOS10), and organelle transport (myosin 19, MYO19 (25)). GO enrichment analysis of the 35 differentially abundant mitochondrial proteins did not reveal any statistically significant overrepresented functional pathways. The mitochondrial abundance of 14 of the 35 proteins (40%) was significantly increased upon infection with wild-type *L. monocytogenes*, whereas it decreased for 8 proteins (23%) (Table S1). Interestingly, 7 of the 14 upregulated proteins and 4 of the 8 downregulated proteins did not display these changes upon infection with LLO-deficient bacteria (Table S1), indicating that LLO triggers such alterations in their mitochondrial abundance.

We further focused our attention on proteins predicted to participate in mitochondrial dynamics or membrane-remodeling processes. Interestingly, we found Mic10 among the seven proteins enriched in mitochondria in response to infection by wild-type but not LLO-deficient *L. monocytogenes* (Figure 1E). Mic10 is a core subunit of the mitochondrial contact site and cristae organizing system (MICOS) complex, a conserved IMM complex responsible for the formation of crista junctions (sites where the IMM invaginates to form cristae) (20). Importantly, Mic10 is a small transmembrane protein, whose V-shaped membrane topology and ability to oligomerize provide it with membrane-bending properties that are fundamental for driving the formation of crista junctions (26, 27).

### Mic10 is required for *L. monocytogenes-induced* mitochondrial network fragmentation

To investigate the role of Mic10 in *L. monocytogenes* infection-induced mitochondrial fission, we either lowered or raised Mic10 levels in host cells before infection and analyzed how the mitochondrial network morphology was affected. To assess the effect of Mic10 depletion, we used U2OS cells because its mitochondrial morphology is better suited for microscopy analysis compared to HCT116 cells. U2OS were transfected with non-targeting control (si-Ctrl) or Mic10-targeting (si-Mic10) siRNAs (Figure 2A), after which they were left uninfected or infected with wild-type or LLO-deficient bacteria. Cells were fixed and mitochondria immunolabeled for confocal microscopy analysis. In agreement with our previous results, si-Ctrl cells showed a typical tubular mitochondrial network, which became fragmented upon infection with wild-type, but not LLO-deficient bacteria (Figure 2B). In si-Mic10 cells, the mitochondrial morphology was similar to that of si-Ctrl cells, indicating that Mic10 knockdown does not affect mitochondrial network shape. However, unlike si-Ctrl cells, mitochondrial fragmentation was not detected in si-Mic10 cells in response to wild-type *L. monocytogenes* (Figure 2B). To obtain an unbiased, quantitative representation of these observations, we used a semi-automated morphometric tool to analyze the mitochondrial network morphology from a large number of cells, which allowed us to calculate a mitochondrial fragmentation degree per cell. Results confirmed that the mitochondrial network of si-Mic10 cells does not undergo fragmentation in the presence of *L. monocytogenes*, in contrast to si-Ctrl cells (Figure 2C). We also performed this experiment in HCT116 cells and obtained similar results (Figure S1), thus corroborating the crucial role of Mic10 in mediating mitochondrial fission in response to *L. monocytogenes* infection.

**Figure 2 –.**
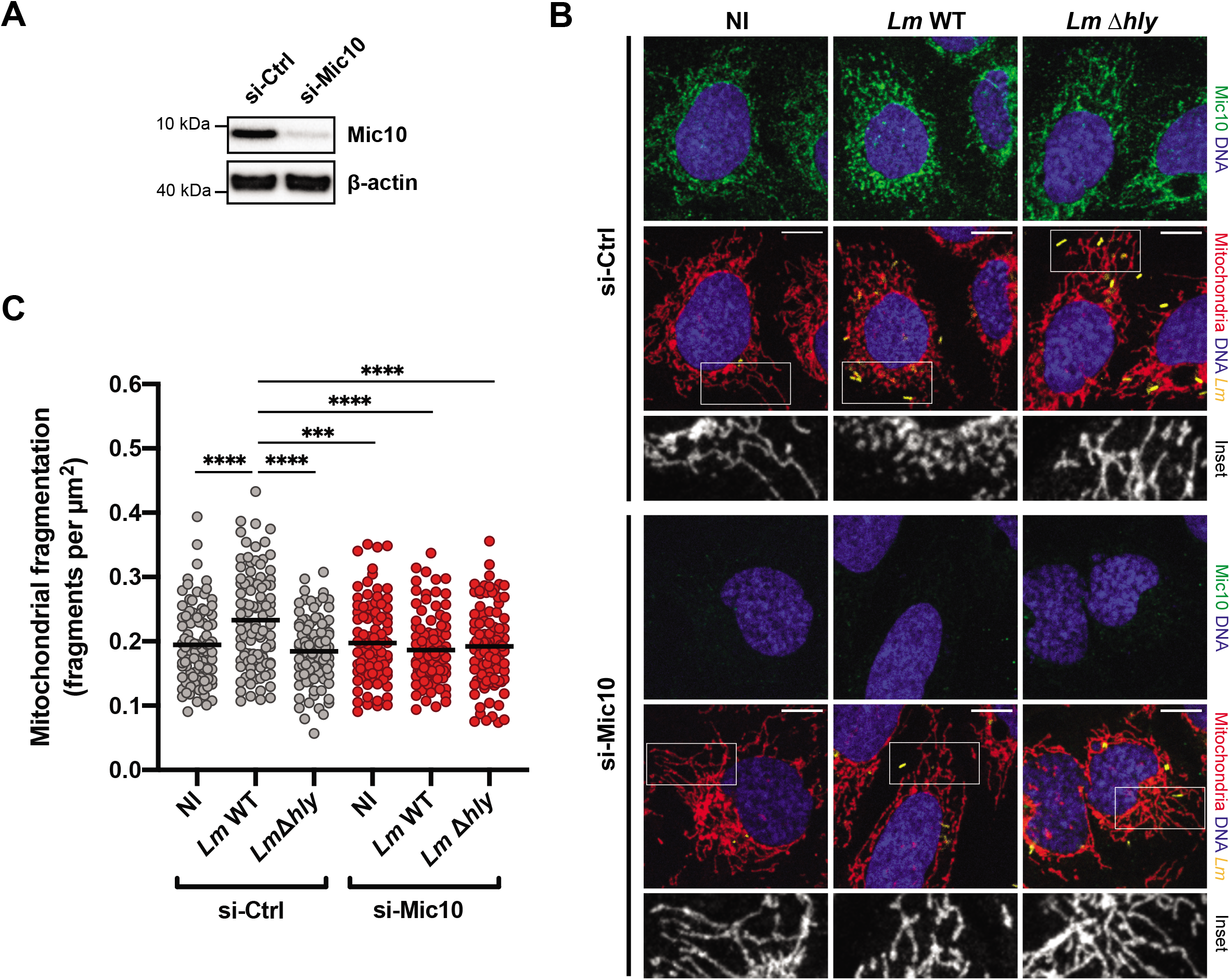
Mic10 knockdown blocks *L. monocytogenes-induced* mitochondrial fragmentation. A) Immunoblot analysis of Mic10 levels in U2OS osteosarcoma cells transfected with control negative (si-Ctrl) or Mic10-targeting (si-Mic10) siRNAs. Beta-actin protein was used as loading control. B) Immunofluorescence analysis of U2OS cells transfected with control negative (si-Ctrl) or Mic10-targeting (si-Mic10) siRNAs, which were left uninfected (NI) or infected (MOI 50, 2h) with GFP-expressing wild type (*Lm* WT) or LLO-deficient (*Lm* Δ*hly*) *L. monocytogenes*. Mic10 is shown in green, mitochondria (anti-Tom20) in red, nuclei (Hoescht 33342) in blue, and bacteria (Lm) in yellow. White box indicates a region of the mitochondrial network magnified (2x) in the inset shown below (mitochondrial labeling only). Scale bar (top right): 10 μm. C) Quantitative analysis of the mitochondrial fragmentation degree in U2OS cells analyzed in B. Mitochondrial network morphology was analyzed using the morphometric ImageJ plugin tool MiNA on mitochondria-labeled images. Fragmentation degree per analyzed cell was determined by the ratio between number of individual mitochondrial particles and total mitochondrial area. Scatter plot graph shows mitochondrial fragmentation degree values for each analyzed cell (dots, n>95) and the mean (horizontal bar). Statistically significant differences were determined by one-way ANOVA with Tukey’s post-hoc test: *** p<0.001; **** p<0.0001.

Next, we examined the effect of increased Mic10 levels in *Listeria*-dependent mitochondrial fission. We followed the experimental approach described earlier but instead of siRNA, cells were transfected with a plasmid driving the constitutive expression of a C-terminal FLAG fusion of the human Mic10 protein (Mic10-FLAG), or the empty parental plasmid as a control. We observed that constitutive expression of Mic10-FLAG resulted in a concomitant reduction of the endogenous Mic10 levels (Figure 3A). This effect was previously reported for both Mic10 and Mic60 and suggests a tight regulation of the total levels of MICOS proteins (28). Whereas control plasmid-transfected cells displayed a fragmented mitochondrial network upon infection with wild-type but not LLO-deficient bacteria (Figure 3B,C), cells expressing high levels of exogenous Mic10-FLAG showed a highly vesiculated mitochondrial network, even in the absence of infection (Figure 3B,C). Unlike Mic10 depletion, which does not affect the mitochondrial network morphology (Figures 2B,C), excessive mitochondrial levels of Mic10 cause a collapse of the mitochondrial network.

**Figure 3 –.**
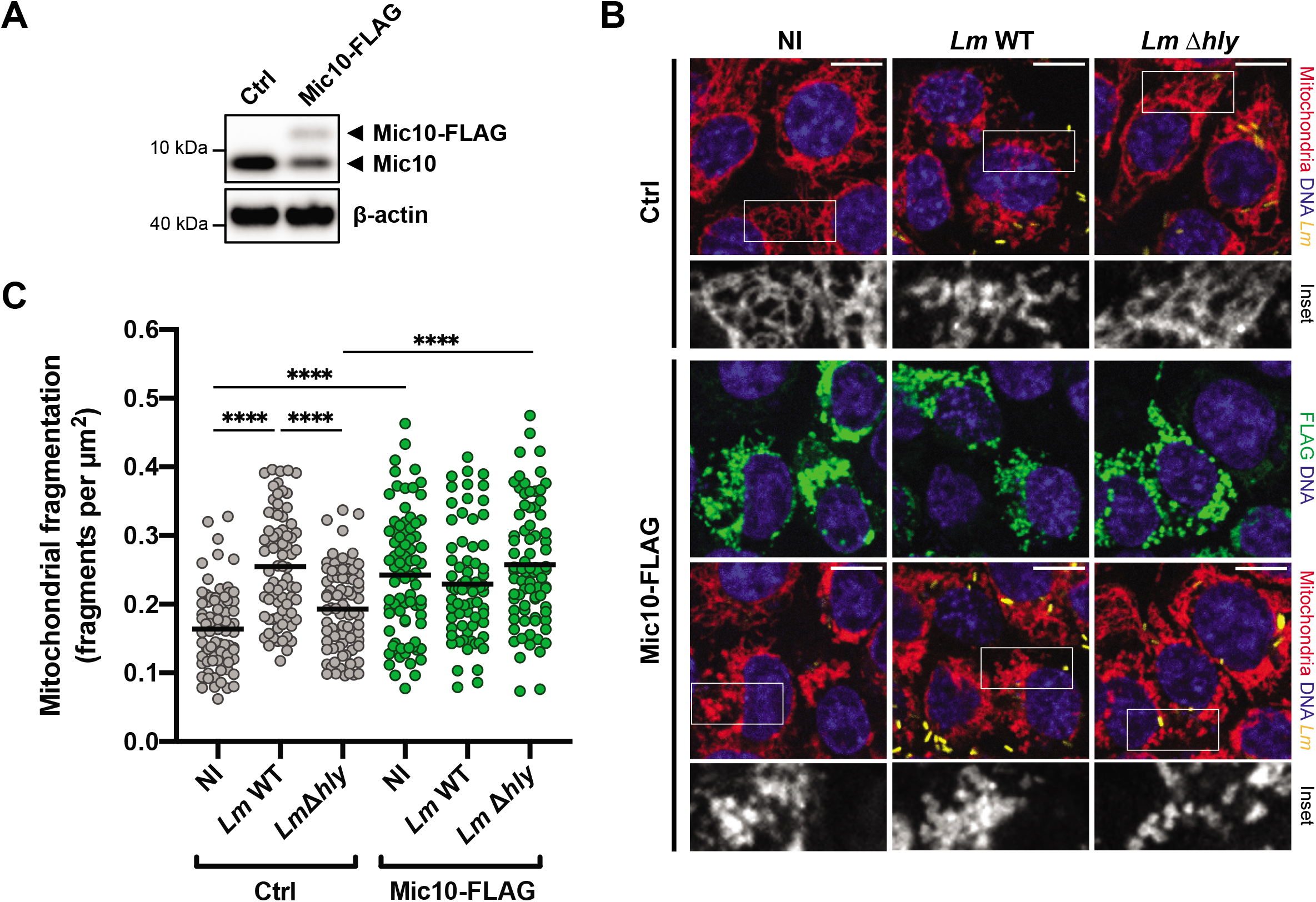
Mic10 overexpression triggers mitochondrial fragmentation regardless of *L. monocytogenes* infection. A) Immunoblot analysis of Mic10 levels in HCT116 cells transiently transfected with control plasmid (Ctrl) or a plasmid constitutively expressing C-terminal FLAG-tagged Mic10 (Mic10-FLAG). Both endogenous Mic10 and exogenous Mic10-FLAG were detected with an anti-Mic10 antibody. Beta-actin protein was used as loading control. B) Immunofluorescence analysis of HCT116 cells transiently transfected with control plasmid (Ctrl) or a plasmid constitutively expressing C-terminal FLAG-tagged Mic10 (Mic10-FLAG), which were left uninfected (NI) or infected (MOI 20, 2h) with GFP-expressing wild type (*Lm* WT) or LLO-deficient (*Lm* Δ*hly*) *L. monocytogenes*. Mitochondria (anti-Tom20) is shown in red, Mic10-FLAG (anti-FLAG) in green, nuclei (Hoescht 33342) in blue, and bacteria (*Lm*) in yellow. White box indicates a region of the mitochondrial network magnified (2x) in the inset shown below (mitochondrial labeling only). Scale bar (top right): 10 μm. C) Quantitative analysis of the mitochondrial fragmentation degree in cells analyzed in B. Mitochondrial network morphology was analyzed using the morphometric ImageJ plugin tool MiNA on mitochondria-labeled images. Fragmentation degree per analyzed cell was determined by the ratio between number of individual mitochondrial particles and total mitochondrial area. Scatter plot graph shows mitochondrial fragmentation degree values for each analyzed cell (dots, n>70) and the mean (horizontal bar), and is representative of three independent experiments. Statistically significant differences were determined by one-way ANOVA with Tukey’s post-hoc test: **** p<0.0001.

These results indicate that *L. monocytogenes* requires basal Mic10 levels to trigger mitochondrial fission in an LLO-dependent manner. Moreover, together with our proteomic data, they suggest that this mitochondrial network breakdown could be a result of increased Mic10 levels in mitochondria.

### Mic10 contributes to an efficient *L. monocytogenes* cellular infection

The dynamic state of the mitochondrial network was reported to play a role in the early steps of *L. monocytogenes* cellular infection, as cells with fragmented mitochondria were less susceptible to infection, whereas cells with hyperfused mitochondria showed improved infection levels (16). Considering our results regarding the effect of Mic10 levels on the morphological status of the mitochondrial network, we wondered whether and how Mic10 levels affect *L. monocytogenes* infection. We performed gentamicin protection assays in control cells and in cells either depleted of Mic10 or overexpressing Mic10, and after infection with wild-type bacteria, we quantified the intracellular bacterial load.

Compared to control cells, Mic10-depleted cells were 30% less infected (Figure 4A), whereas cells overexpressing Mic10 showed a 20% increase in infection (Figure 4B). To determine if the reduced infection levels observed under Mic10 depletion were due to alterations in cellular bioenergetics elicited by defects in mitochondrial function and energy metabolism, we analyzed the mitochondrial respiratory and ATP production capacity of these cells. The oxygen consumption rate and ATP levels in si-Mic10 cells were similar to those in si-Ctrl cells (Figure 4C,D), in agreement with other studies (29, 30). These results suggest that the effect of Mic10 on *L. monocytogenes* infection is not caused by changes in mitochondrial energy production.

**Figure 4 –.**
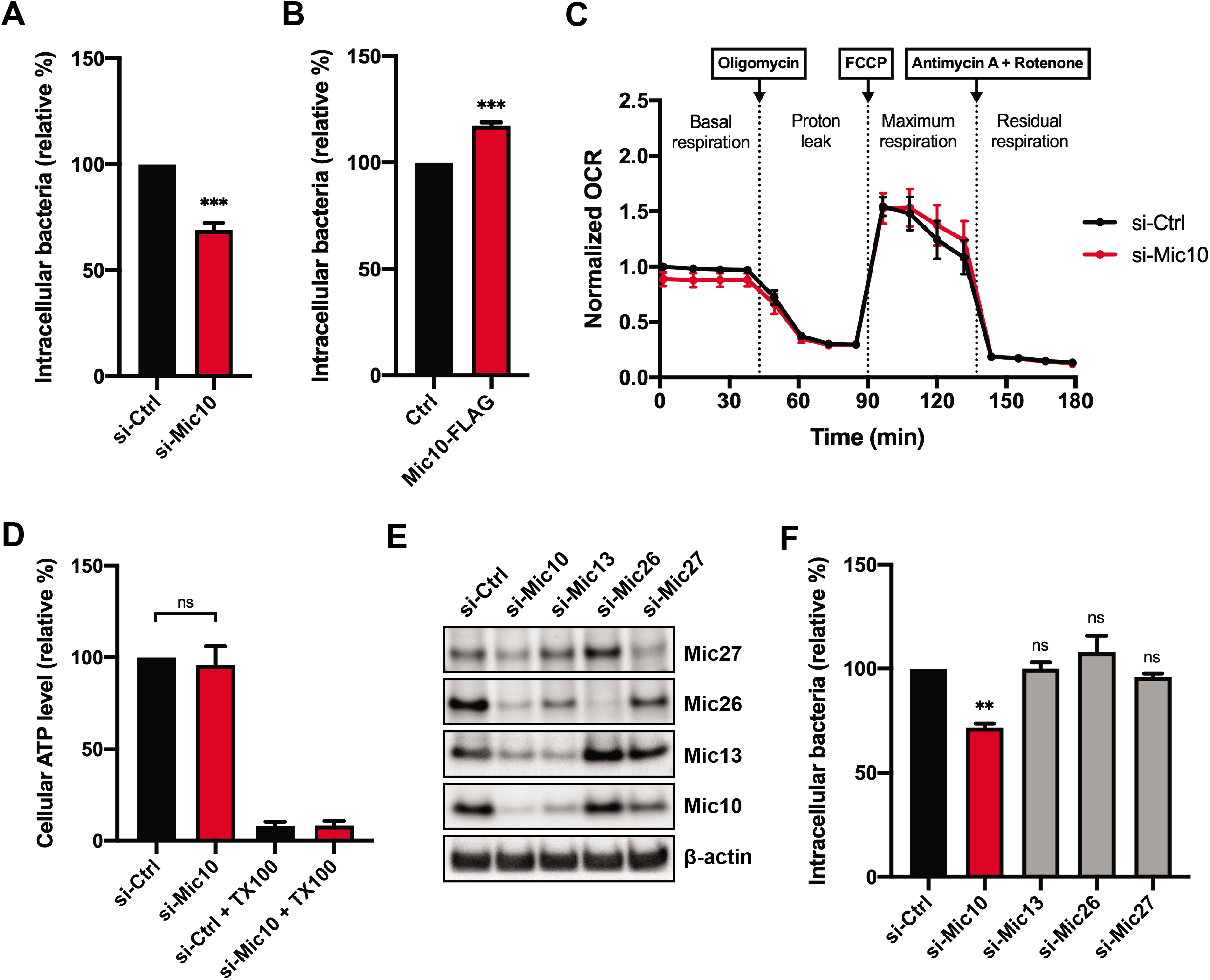
Mic10 contributes to *L. monocytogenes* cellular infection. A) Quantification of intracellular bacteria in HCT116 cells transfected with control negative (si-Ctrl) or Mic10-targeting (si-Mic10) siRNAs, following infection with wild type *L. monocytogenes* (MOI20, 2h). Results are shown as mean ± SEM of three independent experiments, and represented as percentage of intracellular bacteria relative to those quantified in si-Ctrl cells. Statistically significant difference was determined by unpaired, two-tailed t-test: *** p<0.001. B) Quantification of intracellular bacteria in HCT116 cells transiently transfected with control plasmid (Ctrl) or a plasmid constitutively expressing C-terminal FLAG-tagged Mic10 (Mic10-FLAG), following infection with wild type *L. monocytogenes* (MOI 20, 2h). Results are expressed as percentage of intracellular bacteria relative to those quantified in Ctrl cells, and shown as mean ± SEM of three independent experiments. Statistically significant difference was determined by unpaired, two-tailed t-test: *** p<0.001. C) Oxygen consumption rate (OCR, pmol/min) of HCT116 cells transfected with control negative (si-Ctrl) or Mic10-targeting (si-Mic10) siRNAs was measured in a Seahorse XF Analyzer. Electron transport chain inhibitors (oligomycin, FCCP, and antimycin A/rotenone) were added at defined time points to monitor specific components of cellular respiration. Results are expressed as fraction of the first OCR value (basal respiration) of si-Ctrl cells, and shown as mean ± SEM of three independent experiments. D) Cellular ATP levels in HCT116 cells transfected with control negative (si-Ctrl) or Mic10-targeting (si-Mic10) siRNAs were quantified by luminescence-based plate assay, using ATPlite Luminescence Assay kit. Negative controls consist of cells treated with Triton X-100 (TX100). Results are expressed as percentage of ATP levels relative to those quantified in si-Ctrl cells, and shown as mean ± SEM of three independent experiments. Statistically significant differences were determined by unpaired, two-tailed t-test: ns, not significant. E) Immunoblot analysis of the levels of Mic10 subcomplex members (Mic10, Mic13, Mic26 and Mic27) in HCT116 cells transfected with control negative siRNA (si-Ctrl), or siRNA targeting Mic10 (si-Mic10), Mic13 (si-Mic13), Mic26 (si-Mic26) or Mic27 (si-Mic27). Beta-actin protein was used as loading control. F) Quantification of intracellular bacteria in cells treated as in E, after infection with wild type *L. monocytogenes* (MOI 20, 2h). Results are shown as mean ± SEM of three independent experiments, and represented as percentage of intracellular bacteria relative to those quantified in si-Ctrl cells. Statistically significance was determined by one-way ANOVA with Dunnett’s post-hoc test: ns, not significant; ** p<0.01.

Depletion of Mic10 in yeast and mammalian cells results in reduced levels of MICOS proteins Mic13, Mic26 and Mic27 (26, 28, 31, 32), which interact closely with Mic10 to form the Mic10 subcomplex (20). We thus assessed if the decreased *Listeria* infection associated with Mic10 knockdown was a consequence of the lower abundance of Mic13, Mic26 and/or Mic27 by performing siRNA-mediated silencing of the corresponding genes before infection with wild-type *L. monocytogenes*. We confirmed that Mic10 depletion results in partial downregulation of the other three members of the Mic10 subcomplex (Figure 4E). In turn, Mic13 and, to a lesser degree, Mic27, are also necessary to sustain basal Mic10 levels (Figure 4E), in agreement with previous reports (31, 32). Quantification of intracellular bacteria showed again impaired infection of Mic10-depleted cells, but revealed no difference between control cells and cells depleted for either Mic13, Mic26 or Mic27 (Figure 4F). This surprising result demonstrates that none of these MICOS subunits is individually required for *Listeria* cellular infection, and therefore supports a unique role of Mic10 in this process. In agreement with this finding, besides Mic10, none of the other six subunits of the metazoan MICOS complex (Mic13, Mic19, Mic25, Mic26, Mic27, and Mic60) showed significantly changed levels with *L. monocytogenes* infection in our proteomic analysis. We investigated if increased Mic10 levels resulted from infection-driven transcriptional upregulation. Quantitative real-time PCR analysis of RNAs from cells infected or not with wild-type bacteria showed that the relative level of transcripts coding for Mic10 or any other MICOS subunits remained unchanged upon *L. monocytogenes* infection (Figure S2). This result suggests that instead of upregulating Mic10 transcription to elevate its protein levels, *L. monocytogenes* infection promotes an accumulation of Mic10 in mitochondria through an unknown post-transcriptional mechanism.

Overall, these results indicate that *Listeria* cellular infection efficiency is specifically and positively correlated with increased mitochondrial levels of Mic10.

## Discussion

Bacterial pathogens like *L. monocytogenes, Legionella pneumophila, Shigella flexneri, Chlamydia trachomatis* and *Mycobacterium tuberculosis* interfere with mitochondrial dynamics to create an intracellular environment suited for survival and persistence (14–16, 18, 33–35). In particular, *L. monocytogenes* was shown to induce mitochondrial fission early in infection and cause a metabolic slowdown (16, 18), delaying mitochondria-dependent cellular responses, such as type III interferon signaling (36). Here, we employed quantitative proteomics to characterize the mitochondrial response to *L. monocytogenes* infection and search for host factors involved in *L. monocytogenes-induced* mitochondrial fission. We revealed the MICOS complex protein Mic10 as a new player in this process. We showed that *L. monocytogenes* infection increases Mic10 levels in mitochondria in an LLO-dependent manner, and that Mic10 abundance is positively correlated with *L. monocytogenes-induced* mitochondrial fragmentation and host cell infection. This supports a model whereby *L. monocytogenes* infection promotes elevated mitochondrial Mic10 levels to trigger organellar fission and favor cellular infection.

Our proteomic approach yielded a degree of mitochondrial enrichment (36%) and mitochondrial proteome coverage (53%) comparable to those reported in other studies using varied mitochondrial isolation and mass spectrometry protocols (37–39). One of these studies explored the host mitochondrial response to *M. tuberculosis* infection, showing that virulent strains increased mitochondrial energy production and protected host cells from apoptosis, as opposed to avirulent bacteria (37). These changes were partially supported at the protein level, with upregulation of proteins involved in respiration and anti-apoptotic mechanisms, and reduced levels of proteins linked to anti-microbial response. Our proteomic data also hint that mitochondrial translation and respiration are enhanced in response to *L. monocytogenes* infection, possibly to compensate for the drop in mitochondrial membrane potential (16).

We chose to explore Mic10 because it represents a *bona fide* IMM protein with well-characterized membrane-shaping properties (26, 27, 40–42), and its mitochondrial levels showed an LLO-dependent upsurge with infection. We hypothesized that elevated Mic10 levels could drive deregulated IMM remodeling, resulting in mitochondrial fission. In support of this assumption, we showed that *L. monocytogenes* cannot fragment mitochondria in Mic10-depleted cells. In contrast, we observed clear mitochondrial fragmentation in cells overexpressing Mic10, even in the absence of bacteria. Others have reported that excessive Mic10 levels disrupt cristae structure (26), and that Mic10 knockdown or knockout also result in absent cristae junctions and unattached cristae stacked in the matrix (26–30, 40, 43). These phenotypes showcase the importance of Mic10 in IMM structure maintenance and suggest that *L. monocytogenes* could target Mic10 to induce IMM remodeling and trigger mitochondrial fission. We did not observe changes in the mitochondrial morphology of Mic10-depleted cells, implying that the ultrastructural defects caused by Mic10 knockdown are not sufficient to elicit mitochondrial fragmentation, in contrast to the disruptive effect of excessive Mic10 levels. Consistently, knockdown of other MICOS subunits, such as Mic60 (28), Mic19 and Mic25 (44), did not cause mitochondrial fragmentation, although Mic60- and Mic19-depleted mitochondria showed bulb-like enlargements (28, 44). Similar features were reported in Mic10-null yeast mitochondria (45), but we did not observe them in our si-Mic10 cells.

Surprisingly, the enrichment of Mic10 in mitochondria upon *L. monocytogenes* infection was not due to increased Mic10 transcription, which suggests that Mic10 accumulation in mitochondria occurs at the protein level. This could be caused by increased import or reduced turnover of Mic10 in mitochondria. As a nuclear gene-encoded protein, Mic10 is imported from the cytosol via the mitochondrial protein import machinery (46, 47). However, as other MICOS proteins are similarly imported (47), and our proteomics data showed no significant changes in their mitochondrial levels, it seems unlikely that increased Mic10 levels are caused by enhanced mitochondria import. Protein turnover in mitochondria is carried out by multiple proteases residing in the different mitochondrial compartments (48). Interestingly, two of these proteases, Yme1L and Oma1, were reported to participate in the processing of Mic60 and Mic19, respectively (28, 44), suggesting that they may participate in Mic10 proteolysis. Future knockdown or loss-of-function experiments should clarify the involvement of Yme1L and/or Oma1 in Mic10 turnover.

Cells with fragmented mitochondria were shown to be less infected by *L. monocytogenes*, raising the hypothesis that pre-fragmented mitochondria are more resistant to the *L. monocytogenes*-induced bioenergetic slowdown (16). Here, we demonstrate that *L. monocytogenes* infection is partially impaired in cells with reduced Mic10 abundance, suggesting that Mic10-dependent mitochondrial fission induced by *L. monocytogenes* is important for subsequent cellular infection. In contrast, bacterial infection was improved by 20% in cells transfected with DNA driving Mic10 overexpression. Since not every cell overexpressed Mic10, it is possible that this margin may be higher. Further experiments using stable clones of Mic10-overexpressing cells will be helpful to confirm whether *Listeria* infection is enhanced due to increased mitochondrial Mic10 levels.

An important question is how *L. monocytogenes* manipulates events taking place inside mitochondria, even at early steps of infection when it is entering host cells or possibly still in the extracellular medium. The obvious trigger is LLO secreted by *L. monocytogenes*, which mediates Ca^2+^ influx into the host cytoplasm (16, 49, 50). Mitochondria take up Ca^2+^ from the cytosol via the mitochondrial calcium uniporter (MCU) complex (3), which includes the mitochondrial calcium uptake protein 1 (MICU1) that controls the Ca^2+^ concentration crossing the MCU channel (51). Interestingly, MICU1 was identified in our proteomic analysis, showing an apparent enrichment with *L. monocytogenes* infection in an LLO-dependent manner (Table S1). Mitochondrial Ca^2+^ efflux is mediated, among others, by the sodium/calcium exchanger NCLX (52), which can be activated by protein kinase A (PKA)-mediated phosphorylation (53). The catalytic subunit alpha of PKA (PRKACA) is one of four mitochondria-related proteins that are less abundant with infection in an LLO-dependent manner, suggesting that PKA-mediated NCLX activation is impaired during *L. monocytogenes* infection. MICU1 upregulation and NCLX inhibition could result in increased mitochondrial Ca^2+^ concentration and, among other effects, a generalized collapse of the mitochondrial network (54). An investigation on the contribution of these mitochondrial proteins could clarify a role for mitochondrial Ca^2+^ uptake in *L. monocytogenes-* and possibly also Mic10-dependent mitochondrial fragmentation.

In conclusion, this work represents the first proteomic analysis of the mitochondrial response to *L. monocytogenes* infection and allowed us to reveal a novel actor in mitochondrial dynamics, which is specifically manipulated by *L. monocytogenes* to create the ideal setting for host cell infection.

## Materials and methods

### Bacterial strains, cell lines and growth conditions

The following *Listeria monocytogenes* strains were used in this study: wild type EGD (BUG 600), its isogenic LLO mutant EGDΔ*hly* (BUG 3650), and the corresponding GFP-expressing derivatives EGD-cGFP (BUG 2539) and EGDΔ*hly*-cGFP (BUG 2786). Bacteria were grown at 37 °C in brain heart infusion (BHI) media (Difco, BD), supplemented with chloramphenicol (7 μg/mL), when required. The following tissue culture cell lines were used in this study: HCT116 (human colorectal adenocarcinoma; ATCC CCL-247) and U2OS (human osteosarcoma; ATCC HTB-96). Cells were maintained in McCoy’s 5A GlutaMAX medium (Gibco), supplemented with 1 mM non-essential amino acids (Gibco) and 10% (v/v) fetal bovine serum (FBS) (BioWest), and grown at 37 °C in a humidified 10% CO_2_ atmosphere.

### Cell transfection

For transient gene knockdown, cells were reverse transfected with siRNAs in 24-well plates, using Lipofectamine RNAiMax (Invitrogen) according to the manufacturer’s instructions, except that McCoy’s 5A was used as dilution medium. The medium was changed the following day and cells were assayed 48 h post-transfection. siRNA duplexes were used at the following concentrations: siRNA Universal Negative Control #1 (Sigma-Aldrich) and Mic10 (5’-CGGAUGCGGUCGUGAAGAUtt-3’; Eurofins Genomics) at 100 nM; Mic13 (Ambion, Silencer Select #s195661), Mic26 (Ambion, Silencer Select #s35601), and Mic27 (Ambion, Silencer Select #s225655) at 20 nM. For transient overexpression of Mic10, cells were seeded in 24-well plates one day before transfection with 0.5 μg of Mic10-FLAG plasmid DNA (pcDNA3.1(+)-MINOS1-DYK; GenScript, ORF cDNA clone ID OHu15514), using jetPRIME (Polyplus Transfection) according to the manufacturer’s instructions. Control cells were transfected with empty plasmid DNA (pcDNA3.1(+); Invitrogen). Cells were assayed 24 h post-transfection. For immunofluorescence, cells were seeded in wells containing glass coverslips.

### Cell infections

For cell infection assays, confluent monolayers were incubated for 1 h at 37 °C in FBS-free cell culture medium alone (non-infected cells) or inoculated with logarithmic phase bacteria (OD_600nm_ 0.6–1.0) at a multiplicity of infection (MOI, bacteria/cell) of 20 (for HCT116 cells) or 50 (for U2OS cells). Medium was removed and cells were incubated for another hour (total infection time: 2 h) at 37 °C in FBS-containing culture medium supplemented with 20 μg/mL gentamicin sulfate (Sigma-Aldrich), to kill extracellular bacteria. Cells were then washed with Dulbecco’s phosphate-buffered saline (DPBS; Gibco) before processing for further analyses. For immunofluorescence, cells were infected with GFP-expressing *L. monocytogenes* strains. To quantify intracellular bacterial levels, infected cells were lysed in ice-cold 0.2% (v/v) Triton X-100 in DPBS, serially diluted in DPBS, and plated on BHI agar plates. Colony-forming units (CFUs) were counted after 24 h of incubation at 37 °C and bacterial numbers were normalized to the inoculum concentration.

### Mitochondrial isolation and LC-MS/MS sample preparation

For label-free quantitative proteomic analysis of mitochondria, HCT116 cells (~5×10^7^) cultivated in 150-mm dishes were treated as outlined in Figure 1A. Cells were left uninfected (NI) or infected either with wild type (*Lm* WT) or LLO-deficient EGD (*Lm* Δ*hly*), as described above. Three independent biological replicates were prepared and analyzed for each condition. After infection, mitochondria were isolated from cells by magnetic immunoaffinity separation, using the Mitochondria Isolation Kit (human; Miltenyi Biotec). For this, cells were washed with ice-cold DPBS and scraped in ice-cold kit lysis buffer (1 mL/10^7^ cells) containing a cocktail of protease inhibitors (cOmplete, EDTA-free; Roche). Cells were then lysed in a Potter-Elvehjem homogenizer (~50 strokes), with lysis monitored by trypan blue staining. Lysates were centrifuged for 5 min at 800×g (4 °C) to pellet unbroken cells, and the supernatant was recovered for magnetic labeling and separation of mitochondria as detailed in the kit instructions. Purified mitochondria were resuspended in urea lysis buffer (20 mM HEPES pH 8.0, 8 M urea) and protein concentration was measured with BCA Protein Assay kit (Pierce). Proteins were reduced for 30 min at 55 °C in the presence of 25 mM DTT, and then alkylated for 15 min in the dark in the presence of 50 mM iodoacetamide. Samples were diluted two-fold with 20 mM HEPES pH 8.0 and proteins digested with Lys-C (Promega) at a protease/protein ratio of 1:100 (w/w) for 4 h at 37 °C. Samples were diluted two-fold again and incubated overnight at 37 °C with trypsin (sequencing grade modified, Promega) at a 1:50 (w/w) ratio. Formic acid (FA) was added at 1% (v/v), and after 10 min on ice, samples were centrifuged for 10 min at 10,000×g to pellet any insoluble material. Peptides in the supernatant were purified in Sep-Pak C18 cartridges (100 mg; Waters), lyophilized, dissolved in solvent A [0.1% (v/v) FA in water/acetonitrile (ACN) (98:2, v/v)] and quantified by absorbance at 280 nm (NanoDrop, Thermo Fisher Scientific). Samples were analyzed by LC-MS/MS as described in Text S1.

### Immunofluorescence

Cells grown on glass coverslips were fixed for 15 min at room temperature in 4% (v/v) paraformaldehyde in PBS, permeabilized for 5 min in 0.5% (v/v) Triton X-100 in PBS, and blocked for 20 min in blocking buffer [1% (w/v) BSA, 10% (v/v) goat serum in PBS]. Labelling with primary and fluorophore-conjugated secondary antibodies or dyes was performed in blocking buffer for 1 h at room temperature. Cells were washed three times in PBS between every step after fixation, except after blocking. Coverslips were mounted onto microscope slides with FluoroMount-G mounting medium (Interchim), and imaged the next day or stored in the dark at 4 °C. Primary antibodies were used as follows: rabbit polyclonal anti-C1ORF151/Mic10 (1:200; Abcam, ab84969), mouse monoclonal anti-Tom20 clone 29 (1:200; BD Transduction Laboratories), rabbit polyclonal anti-Tom20 clone F-10 (1:200; Santa Cruz Biotechnology), and mouse monoclonal anti-FLAG clone M2 (1:100; Sigma-Aldrich). Anti-rabbit and anti-mouse antibodies conjugated to Alexa Fluor 568 and 647 dyes (1:500; Molecular Probes) were used as secondary antibodies; Hoescht 33342 (Molecular Probes) was used to stain DNA. Cells were analyzed in a ZEISS AxioObserver.Z1 inverted microscope (Carl Zeiss AG) equipped with a high-speed CSU-X1 spinning-disk confocal system (Yokogawa) and an Evolve EM-CCD camera (Photometrics). Single focal plane images were acquired through a Plan-Apochromat 63×/1.4 Ph3 oil objective across multiple wavelength channels, using MetaMorph software (version 7.7.9.0). Fiji was used for image processing, including channel color selection, brightness and contrast adjustment, addition of scale bars and generation of composite images.

### Mitochondrial morphology analysis

Confocal images of cells were taken from various fields of view randomly selected across the entire coverslip area and their mitochondrial morphology was analyzed using the semi-automated morphometric tool MiNA within Fiji (55). Mitochondrial networks (labeled with anti-Tom20) from individual cells were selected and digitally isolated before batch analysis. From the output data, a ratio of the values listed under the “Individuals” (number of unbranched mitochondrial particles, e.g. puncta and rods) and “Mitochondrial footprint” (mitochondrial area) parameters was calculated to determine the degree of mitochondrial fragmentation per analyzed cell. A minimum of 50 cells were analyzed per condition, in a total of three independent experiments.

### Statistics

Statistical analyses were performed in Prism 8 (GraphPad Software). Unpaired two-tailed Student’s t-test was used to compare the means of two groups; one-way ANOVA was used with Tukey’s post-hoc test for pairwise comparison of means from more than two groups, or with Dunnett’s post-hoc test for comparison of means relative to the mean of a control group. Difference between group means were considered statistically significant at p-value < 0.05. Significance levels are indicated as: ns, not significant (p > 0.05); *, p < 0.05; **, p < 0.01; ***, p < 0.001, ****, p < 0.0001.

### Data availability

Mass spectrometry proteomics data have been deposited to the ProteomeXchange Consortium via the PRIDE (56) partner repository with the dataset identifier PXD014667.

## Acknowledgements

We thank current and past lab members for helpful discussions; Francis Impens and Evy Timmerman (VIB Proteomics Core, University of Ghent, Belgium) for training and assistance with proteomic analyses; and Alessandro Pagliuso for critical reading of the manuscript.

This study was supported by grants to P.C. from the European Research Council (H2020-ERC-2014-ADG 670823-BacCellEpi), the Agence Nationale de la Recherche (ANR) and the French Government’s “Investissements d’Avenir” program Laboratoires d’Excellence “Integrative Biology of Emerging Infectious Diseases” (LabEx IBEID, ANR-10-LABX-62-IBEID). A.S. was supported by a BioSPC doctoral fellowship from the Université Paris Diderot. P.C. is a Senior International Research Scholar of the Howard Hughes Medical Institute. F.S. is a CNRS permanent researcher.

---

**Figure S1 – Confirmation of Mic10-dependent *L. monocytogenes*-induced mitochondrial fragmentation in HCT116 cells.**

A) Immunoblot analysis of Mic10 levels in HCT116 cells transfected with control negative (si-Ctrl) or Mic10-targeting (si-Mic10) siRNAs. Beta-actin protein was used as loading control.

B) Immunofluorescence analysis of HCT116 cells transfected with control negative (si-Ctrl) or Mic10-targeting (si-Mic10) siRNAs, which were left uninfected (NI) or infected (MOI 20, 2h) with GFP-expressing wild type (*Lm* WT) or LLO-deficient (*Lm* Δ*hly*) *L. monocytogenes*. Mic10 is shown in green, mitochondria (anti-Tom20) in red, nuclei (Hoescht 33342) in blue, and bacteria (Lm) in yellow. White box indicates a region of the mitochondrial network magnified (2x) in the inset shown below (mitochondrial labeling only). Scale bar (top right): 10 μm.

C) Quantitative analysis of the mitochondrial fragmentation degree in cells analyzed in B. Mitochondrial network morphology was analyzed using the morphometric ImageJ plugin tool MiNA on mitochondria-labeled images. Fragmentation degree per analyzed cell was determined by the ratio between number of individual mitochondrial particles and total mitochondrial area. Scatter plot graph shows mitochondrial fragmentation degree values for each analyzed cell (dots, n>50) and the mean (horizontal bar), and is representative of three independent experiments. Statistically significant differences were determined by one-way ANOVA with Tukey’s post-hoc test: * p<0.05; ** p<0.01, *** p<0.001.

**Figure S2 – Transcription of Mic10 or any other MICOS complex genes is not upregulated upon *L. monocytogenes* infection.** Analysis of gene expression of human MICOS complex subunits Mic10, Mic13, Mic19, Mic25, Mic26, Mic27, and Mic60 in response to *L. monocytogenes* infection. HCT116 cells were left uninfected (NI) or infected (MOI 20, 2h) with wild type (*Lm* WT) or LLO-deficient (*Lm* Δ*hly*) *L. monocytogenes*, and total cellular RNAs were isolated and used for RT-qPCR analysis. Results are shown as mean ± SEM of three independent experiments, and represented as fold change in transcript levels relative to those in NI cells.

